# Transcriptomic Response to Neuromuscular Electrical Stimulation in Muscle, Brain and Plasma EVs in WT and Klotho-deficient Mice

**DOI:** 10.1101/2025.07.25.666815

**Authors:** Catherine Anne Cavanaugh, Amanda E. Moore, Nicholas Francis Fitz, Iliya Lefterov, Radosveta Koldamova

## Abstract

Neuromuscular electrical stimulation (NMES) was shown to improve motor activities and daily living. Prior studies indicated extracellular vesicles (EVs) play a role in cellular communication. Here, we evaluated transcriptomic profiles of tibialis anterior muscle, brain, and plasma-derived EVs following NMES of WT and Klotho heterozygous (Kl^HET^). Muscle RNA-seq data demonstrated that in both genotypes the most upregulated functional categories were related to glucose metabolism and response to insulin with pathways uniquely affected in each genotype. There was a similarity of non-coding RNA transcriptome of plasma EVs with functional patterns suggesting response to oxygen and insulin, and long-term synaptic potentiation. Brain transcriptome showed little functional overlap between WT and Kl^HET^ mice. In WT, brain upregulation of genes were related to blood flow and cell adhesion processes while Kl^HET^ shows upregulation of immune function. Results indicate that similar metabolic function is impacted in the location of stimulation, but the distal impact of stimulation on the brain is associated with Klotho deficiency.

## Introduction

Neuromuscular electrical stimulation of muscle contractile activity (NMES) is a noninvasive approach aimed to improve activities of daily living and functional motor ability in elderly and post stroke (Figure 1A) [1,2]. Beneficial for muscular angiogenesis, muscular tetanic specific force, cognitive function, and increasing glycemic control, NMES serves as a supplement to therapeutic recovery efforts and a treatment option with benefit in individuals with metabolic syndrome and venous disease [1,3–5].

**Figure 1.**
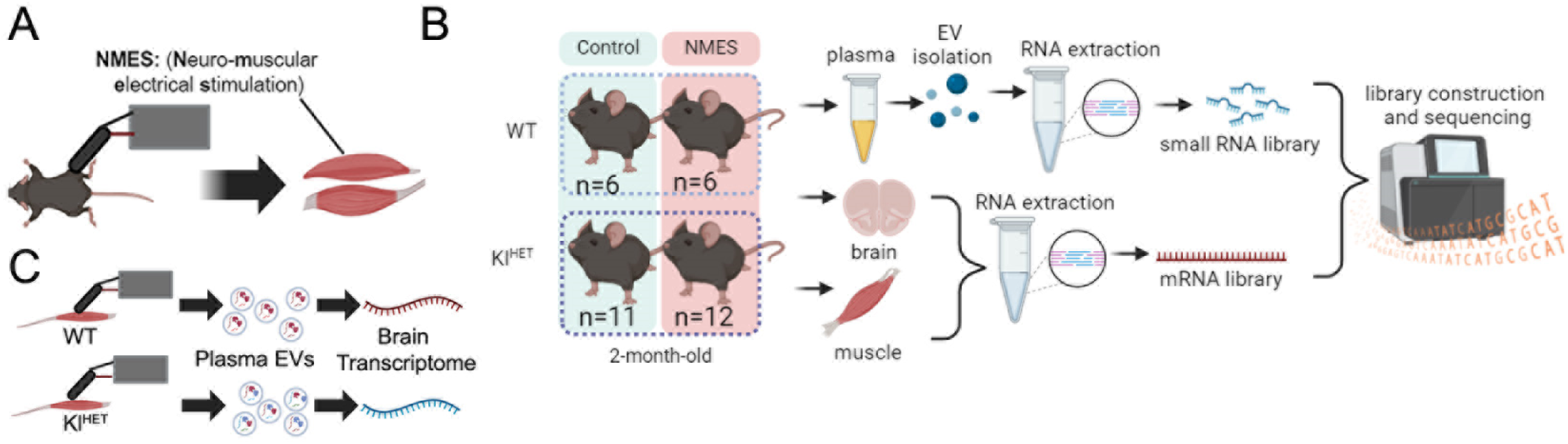
Study design. **(A)** 2-month-old WT and Kl^HET^ mice were subjected to NMES on the tibialis anterior and extensor digitorum longus of each leg for two sets of 10 stimulations every other day for five total sessions. **(B)** Experimental groups were assigned, and tissue was collected. Library generation and sequencing occurred with mRNA libraries from the muscle and brain and small RNA libraries from the plasma-derived extracellular vesicles (EVs). **(C)** The pictorial experimental results summary indicates similarity in transcriptomic response in the area of stimulation and in EVs with the results of the brain transcriptome varying. (N: WT 6/group; Kl^HET^ control=11; Kl^HET^ NMES=12). Figures were made using Biorender.

Emerging information regarding communication between the muscle and brain shows that proteins produced by differentially expressed genes (DEGs), likely carried from the stimulation source by extracellular vesicles (EVs), travel through the bloodstream to the destination tissue, allowing for tissue-crosstalk [6]. Additional support of this work finds similar DEGs from muscle tissue post-direct electrical stimulation and post-aerobic exercise, indicating similar localized benefits and genes with functions such as cell migration, regulating apoptosis, and cell attachment are downregulated in muscle cases of NMES as well as aerobic exercise compared to controls while glucose metabolism is upregulated [7]. Further studies indicate that mRNAs and ncRNAs carried by the EVs can regulate functions within the tissue leading to physiological benefits post-exercise [8]. Therefore, in two of our previous studies we utilized heterochronic blood exchange and tail vein EV injections to better understand the role of circulating neuroprotective young EVs in improving cognitive function [9,10]. Considering the importance of the EVs’ contents-proteins, lipids, ncRNA, and mRNA-being transported between tissues, greater understanding of how DEGs post-exercise regulate the vesicle contents is an important research question. A hindrance in understanding this process comes from the lack of knowledge of how EVs are crossing the blood-brain barrier (BBB) or cerebrospinal fluid (CSF) barrier, which is a future direction [11].

Exerkines are a class of signaling molecules stimulated by exercise that activate the autocrine, paracrine, and/or endocrine systems [12]. The *Klotho* gene encodes the KLOTHO protein, which is expressed in skeletal muscle, kidney, and the choroid plexus, as well as becoming soluble in plasma when cleaved from the membrane via BACE1, ADAM-10 or ADAM-17 [13]. Because *Klotho*-deficient mouse models display reduced lifespan and cognition while *Klotho*-overexpressing mice live longer, *Klotho* is a focus of various fields of research, including neurodegenerative, musculoskeletal, and aging [13–15]. One human participant study noted that modifying diet only compared to diet and exercise treatments noticed that not only was the change in KLOTHO related to an exercise condition, but the concentration of KLOTHO was also related directly to the reduction in weight, fat loss, and waist circumference in patients [16]. Additionally, a study comparing treatments following coronary artery bypass grafting (CABG) noted that greater higher KLOTHO levels following treatment procedures were correlated with improved delayed memory test results [17]. While functional connectivity was seen in NMES rehabilitation patients, there was no significant KLOTHO increase in this condition [17]. These findings have led us to further explore the metabolic interplay between NMES treatment and *Klotho* deficiency to better understand potential transcriptomic effects of noninvasive metabolic treatments.

This study evaluated how NMES treatment affected muscle and brain transcriptome as well as non-coding RNAs expression in plasma EVs (Figure 1B). To determine the role of Klotho deficiency on these parameters, we used previously established NMES protocols to compare the NMES effect in wild type (WT) and klotho-deficient (Kl^HET^) mice [18]. We performed bulk RNA-seq on brain and muscle tissue to determine the effect of treatment and genotype. We showed that in muscle mRNA and plasma-derived EV ncRNA both genotypes included functional overlap related to insulin response (Figure 1C). However, the brain transcriptome showed little functional overlap between WT and Kl^HET^ mice with WT demonstrating upregulation of genes were related to blood flow and cell adhesion processes while Kl^HET^ shows upregulation of immune function (Figure 1C). This experiment works to elucidate the role of localized stimulation and metabolic impact on the brain via EV mRNA and ncRNA trafficking.

## Results

### NMES treatment activates metabolic response in muscle tissue similarly in WT and Kl^HET^ mice

Muscle gene expression profiling was performed by RNA-seq on dissected muscle from WT and Kl^HET^ mice that underwent NMES treatment and edgeR was utilized to identify differentially expressed genes between treatment groups. Our first goal was to determine the effect of NMES, and we compared muscle-derived mRNA from all NMES treated mice to control mice (Figure 2). Volcano plot on Fig. 2A demonstrates the effect of NMES vs control in all mice including WT and Kl^HET^. There were 215 DEGs, 114 up-and 101 downregulated. As visible from the heatmap on Fig. 2B, top up and downregulated DEGs are affected in the same direction in both genotypes. Top biological processes upregulated by all NMES are response to insulin (*Retn, Lep*), positive regulation of insulin secretion (*Ghrl* and *Gck*), response to glucose (*Adipoq*), lipid metabolic process (*Plin1* and *Elovl6*), sarcomere organization (*Mybph*, *Myh6*, and *Myom2*), and gluconeogenesis (*Rbp4* and *Atf3*) (Figure 2.C, upper panel). Figure 2.C (lower panel) illustrates downregulated functions in all NMES versus all control include cell adhesion (*Cyfip2* and *Cdh5*), extracellular matrix organization (*Col12a1*), and cardiac muscle contraction (*Kcnh2* and *Actc1*). These specific GO terms indicate a pattern of modified metabolic function in relation to NMES treatment conditions shown on Figure 2.C. The bar graph on Fig. 2D notes fold changes for selected upregulated DEGs from Fig. 2A related to metabolic processes.

**Figure 2.**
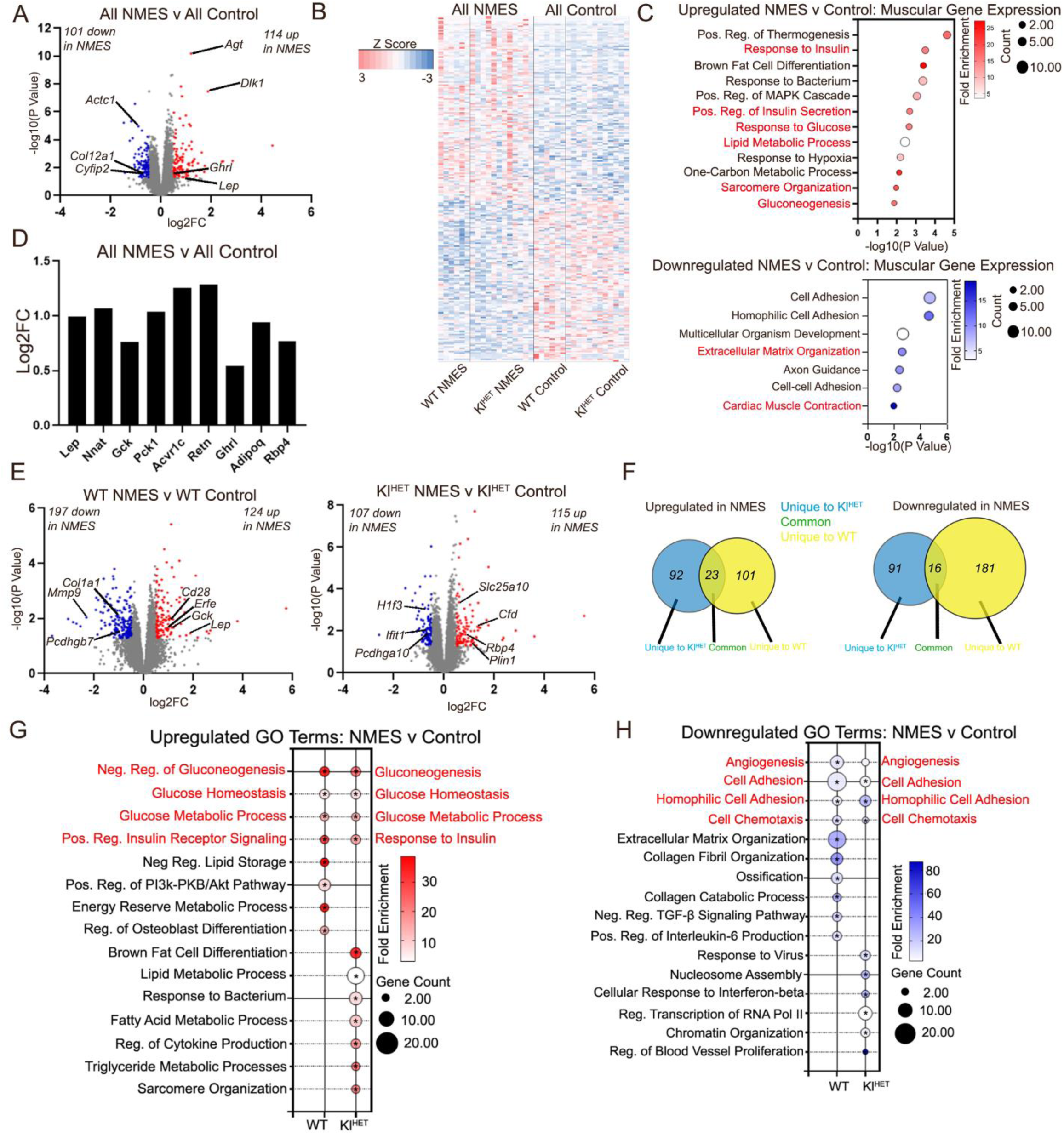
NMES treatment activates metabolic response in muscle tissue similarly in WT and Kl^HET^ **mice**. Gene expression profiling was performed by RNA-seq on dissected muscle from WT and Kl^HET^ mice that underwent NMES treatment and edgeR was utilized to identify differentially expressed genes between treatment groups. **(A)** Volcano plot represents differentially expressed genes (DEG) between NMES and control mice at a cutoff *p*<0.05 and FC>0.5. Red denotes significantly upregulated genes, blue for significantly downregulated, and grey denotes non-significance. **(B)** A heatmap of DEGs. **(C)** Bubble plots depicting Gene Ontology (GO) terms for biological functions associated with DEGs shown on panel A was performed using DAVID. Upregulated terms are shown on the upper panel and downregulated terms on the lower panel. **(D)** Bar graph represents a comparison of fold change of selected DEGs in WT and Kl^HET^. **(E)** Volcano plots of DEGs between NMES and controls in WT (left) and Kl^HET^ (right). **(F)** Venn Diagram represents common and unique DEGs in WT and Kl^HET^ mice as shown on panel E. **(G)** Bubble plots depicting GO terms with the highest fold enrichment for biological functions performed using DAVID. Upregulated are shown on the right panel and downregulated on the left. panel. (N: WT 6/group; Kl^HET^ control=11; Kl^HET^ NMES=12), \**p*<0.05

We next compared how each genotype responds to the treatment. The left panel volcano plot in Fig. 2E illustrates DEGs in WT NMES versus WT control and when compared to DEGs from the Kl^HET^ volcano plot in Fig. 2E, right there is a fairly small number of DEGs shared as a result of treatment (23 common up- and 16 common downregulated genes), emphasizing the necessity to analyze genotype-stratified GO terms (Figure 2.F). In WT, the top upregulated GO term categories include regulation of gluconeogenesis (*Gck*), regulation of phosphatidylinositol 3-kinase/protein kinase B signal transduction (*Cd28*), glucose homeostasis (*Lep*) and regulation of insulin receptor signaling pathway (*Erfe* and *Agt*), and negative regulation of lipid storage (*Lep*) (Figure 2G). For Kl^HET^ mice, top upregulated GO terms are brown fat cell differentiation, lipid metabolic process (*Fasn* and *Plin1*), response to bacterium (*Cfd*), gluconeogenesis (*Slc25a10, Atf3*) and response to insulin (*Rbp4* and *Acvr1c*). Notably, GO terms are shared between the upregulation in WT and Kl^HET^ NMES such as insulin signaling, gluconeogenesis and glucose homeostasis, but the genes included in each GO term are not entirely overlapping. For example, *Erfe*/erythroferrone which is highly expressed in the skeletal muscle and is a part of insulin signaling and gluconeogenesis was differentially upregulated by NMES only in WT but not in Kl^HET^ mice. Previously, *Erfe* was reported to increase after physical training in athletes and acts together with erythropoietin (*Epo*) [19]. However, given a GO term category with similar function upregulated in in Kl^HET^ NMES, response to insulin, genes such as *Retn* which is elevated in subjects with visceral obesity and insulin resistance and *Pck1* which inhibits gluconeogenesis [20, 21]. Overall, the shared upregulated GO terms between the two genotypes as a result of NMES (gluconeogenesis, glucose homeostasis, and insulin response) are all related to metabolic function, emphasizing the relationship between localized muscular treatment and transcriptomic metabolic benefit.

Interestingly, the data indicate less overlap from downregulated DEGs in Fig 2E than compared to upregulated DEGs. Top downregulated GO terms in WT NMES conditions include extracellular matrix organization, cell adhesion, and collagen fibril organization and both categories included numerous collagen genes such as *Col1a1 and Col28a1* and metalloproteases such as *Mmp9* (Figure 2H) [22, 23]. While cell adhesion is downregulated in both WT and Kl^HET^, the Kl^HET^ genes within the category were predominantly protocadherin genes such as *Pcdhga10* which are associated with muscle frailty in older adults [24]. Similarly, homophilic cell adhesion which is a top downregulated term in Kl^HET^ but is expressed in both WT and Kl^HET^ and contains protocadherins in both genotypes. However, none of the genes overlap, with WT genes including *Pcdh17* and *Pcdhgb7* and Kl^HET^ including *Pcdhga10* and *Pcdhga6*. This provides further support that while gene function is similar, the specific DEGs vary which indicates a potentially genotype-mediated transcriptomic response to localized exercise. Other top downregulated GO terms in Kl^HET^ NMES include response to virus and cellular response to interferon-beta (both sharing *Ifit1 and Ifit3*), nucleosome assembly, and RNA transcription of RNA Pol II (*H1f4 and H1f3*).

### KLOTHO deficiency significantly affects muscle transcriptome

Comparisons by genotype examined through muscle gene expression profiling as used in Fig.2 identified DEGs between genotype groups to indicate the role of klotho deficiency in skeletal muscle (Figure 3.A). The volcano plot on Fig. 3A upper panel shows 1077 up- and 282 downregulated DEGs when comparing NMES treatment by genotype including genes such as *Aimp2* which is likely involved in pathogenesis of Parkinson’s disease and related to protein synthesis and cell death [25]. Additionally, the lower panel in Fig. 3A notes 525 upregulated genes such as *Lama3* which is involved in skeletal muscle structural organization and *Pck1* which is involved in both carbohydrate and lipid metabolism and 124 downregulated DEGs when comparing control conditions by genotype [26, 27]. To determine similarities between gene expression in Kl^HET^ and WT genotypes, we created a heatmap of DEGs from Fig. 3A finding that most top upregulated and downregulated genes are impacted in the same direction for each genotype, regardless of treatment (Figure 3.B). Given the large number of upregulated DEGs, REVIGO was used for visualization of GO Terms while bubble plots remained a better representation of GO terms for downregulation. Genotype comparisons within NMES treatment show upregulation of muscular cell processes (*Col18a1*), cellular organization (*Csf1r* and *Itgb4*), and connective tissue development (*Myo5a* and *Col18a1*) and a downregulation of translation (*Aimp2* and *Rpl32*) and transcriptional processes (*Myc*) (Figure 3.C). In Fig. 3.D, control condition comparisons by genotype revealed upregulation in immune modification (*Ifit1*), lipid metabolism (*Pck1* and *Lrrc8c*), and coronary muscular development (*Lrp1*) while translation (*Rpl3*) and transmembrane transport (*Cox7a1*) were downregulated. The venn diagram illustrates a larger overlap in common upregulated DEGs than downregulated DEGs (Figure 3.E) and provides support for the greater GO term overlap in upregulated terms between the NMES and control conditions than downregulation as shown in Fig. 3F.

**Figure 3.**
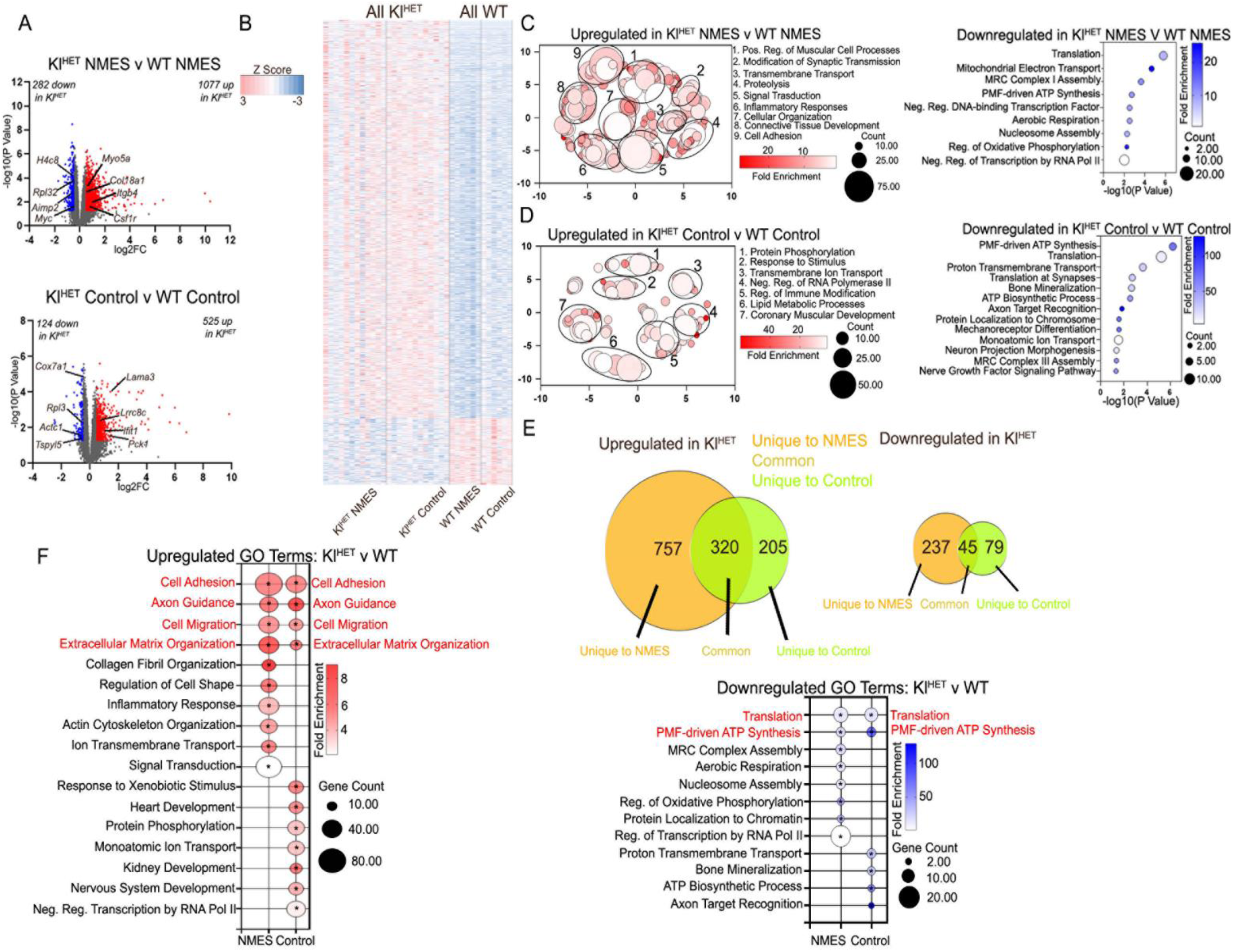
KLOTHO deficiency significantly affects muscle transcriptome. Gene expression profiling was performed by RNA-seq on dissected muscle from WT and Kl^HET^ mice that underwent NMES treatment and edgeR was utilized to identify differentially expressed genes between treatment groups. **(A)** Volcano plots represents differentially expressed genes (DEG) between Kl^HET^ NMES and WT NMES (upper panel) and Kl^HET^ control and WT control (lower panel) mice at a cutoff *p*<0.05 and FC>0.5. Red denotes significantly upregulated genes, blue for significantly downregulated, and grey denotes non-significance. **(B)** A heatmap of DEGs. **(C)** REVIGO for upregulated DEGS (left panel) and bubble plot for downregulated DEGs (right panel) depicting Gene Ontology (GO) terms for biological functions associated with DEGs shown on upper panel A was performed using DAVID. **(D)** REVIGO for upregulated DEGS (left panel) and bubble plot for downregulated DEGs (right panel) depicting Gene Ontology (GO) terms for biological functions associated with DEGs shown on lower panel A was performed using DAVID. **(E)** Venn Diagram represents common and unique DEGs in NMES and control mice as shown on panel A. **(F)** Bubble plots depicting GO terms with the highest fold enrichment for biological functions performed using DAVID. Upregulated are shown on the right panel and downregulated on the left. panel. (N: WT 6/group; Kl^HET^ control=11; Kl^HET^ NMES=12), \**p*<0.05

Notably, shared upregulated GO terms related to structural function include cell adhesion, cell migration, and extracellular organization, all of which share a substantial percentage of genes when comparing exercise conditions. This is likely due to genotype’s impact on animal genome. However, collagen fibril organization and actin cytoskeleton organization are only upregulated in the Kl^HET^ versus WT in the NMES treatment condition which indicates a potential structural increase in muscle that is a result specific to KLOTHO-deficient mice from NMES exercise treatment (Figure 3.F, left). The downregulated GO term overlap is minimal and only includes processes related to translation and proton motive force driven ATP synthesis (Figure 3.F, right).

### Functional patterns emerge in RNA signatures of plasma EVs associated with NMES in WT and Kl^HET^ mice

To better understand how NMES treatment affects plasma EV non-coding transcriptomics, we isolated EV from plasma after NMES and generated small non-codding RNA-seq libraries (ncRNA)s. EV characterization indicated successful isolation and no significant size and concentration difference between genotypes as well as exercise conditions (Figure S.1). Alignment data from both treatments indicate that detected ncRNA readability is similar between NMES and control conditions (Figure 4.A). As with the muscle, we first determined the effect of treatment combining both genotypes. The heatmap shown on Fig. 4B, illustrates differentially enriched miRNA for all NMES versus all control conditions and shows similar directionality between WT and Kl^HET^ comparing all NMES and all control samples. Using DEG miRNA from Fig 4B, DAVID GO Term analysis was conducted finding that top regulated processes likely modified by EV contents include long-term synaptic potentiation, cellular response to lipopolysaccharide, and cellular response to amino acid stimulus (Figure 4.C). Then, miRNA from Fig 4B was aligned to potential gene matches using miRDB to identify GO Term categories of top regulated processes. A STRING diagram was created utilizing the aligned genes detected that fell within the representative GO Term “insulin receptor signaling” (Figure 4.D). Genes such as *Insr* and *Igf1r* which directly modify metabolic and immune function were noted [28]. We then explored how WT and Klotho mice reacted individually to the treatment by comparing NMES group to control in each genotype. Despite the minimal overlap in miRNAs beyond let-7d-5p which contributes to inflammatory responses (Figure 4.E), a larger pattern of functional overlap emerged in Fig 4F [29]. Fig. 4F indicates overlap in functional categories associated with differentially enriched miRNAs such as response to oxygen levels and long-term synaptic potentiation.

**Figure 4.**
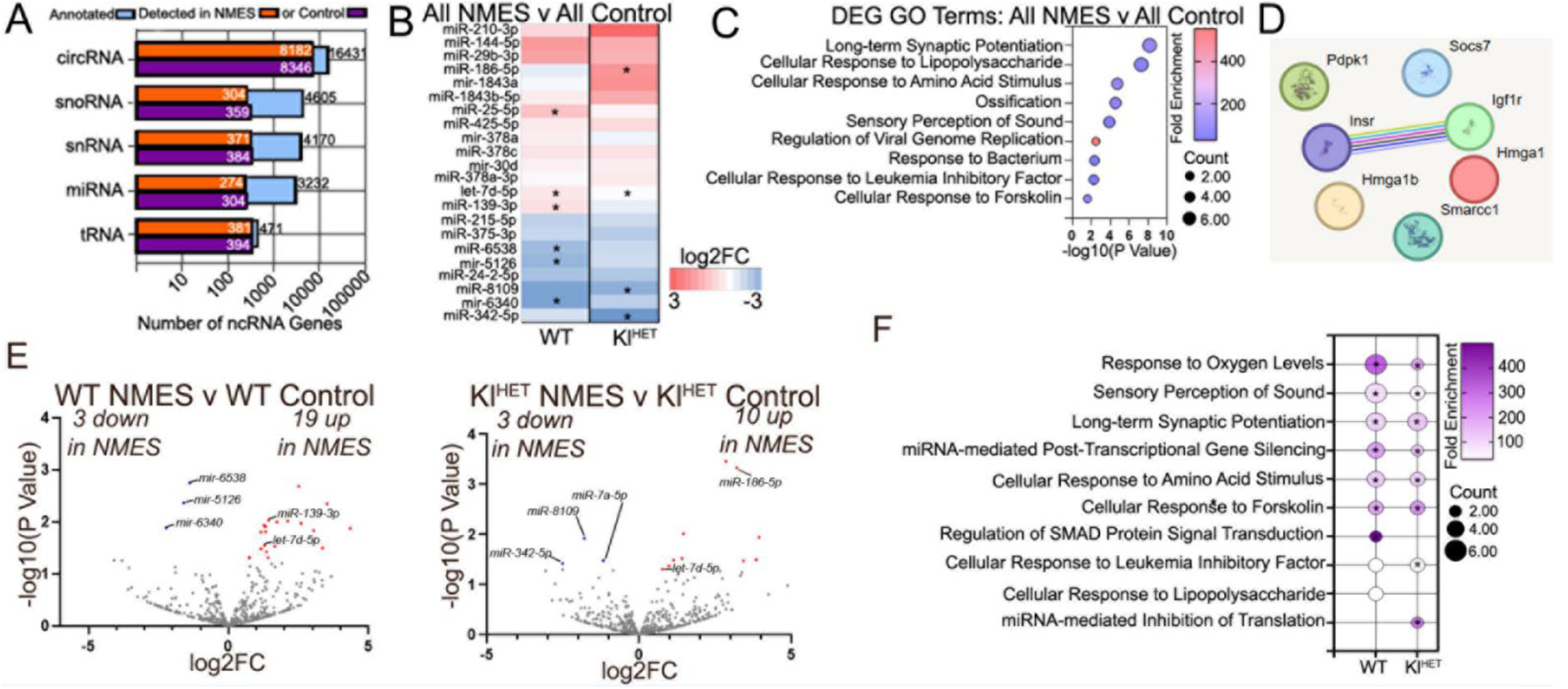
Functional patterns emerge in RNA signatures of plasma EVs associated with NMES in WT and Kl^HET^ mice. RNA was isolated from plasma EVs of WT and Kl^HET^ mice after NMES followed by small noncoding RNA sequencing to determine changes in EV cargos associated with experimental groups. **(A)** Bar graphs showing the alignment for 5 types of noncoding RNAs detected in EV samples (circRNA, snoRNA, snRNA, miRNA, and tRNA) selected for alignment with number of annotations in each treatment group. **(B)** Heat map of fold change in miRNA between all NMES vs all control demonstrates the similarity of the effect in both genotypes. NMES and control include the respective mice from both genotypes. **(C)** GO terms from DEGs miRNAs shown in panel B are visualized. **(D)** STRING interactome for predicted miRNA targeted genes in skeletal muscle GO terms. **(E)** Volcano plots of differentially enriched miRNAs between NMES and control in WT (left) and Kl^HET^ (right). **(F)** GO terms from miRNAs in panel E are visualized. (N: WT 6/group; Kl^HET^ 8/group). \**p*<0.05

### Effect of NMES on brain transcriptome indicates little overlap in WT compared to KlHET

To evaluate the NMES impact on brain transcriptome on WT and Kl^HET^ animals, we performed RNA-seq on whole brain tissue and used edgeR to identify DEGs between treatment groups (Figure 5.A). Volcano plots for WT NMES versus WT control in the left panel of Fig. 5A and Kl^HET^ NMES versus Kl^HET^ control in the right panel of Fig. 4A exhibit the differences of DEGs from NMES treatment (Figure 5.A). In WT mice, the treatment increased the expression of genes such as *Oxt, Avp, Erg, Chrm5,* and *Irs4* which are involved in metabolic function, blood flow, and synaptic plasticity [30–32] (Figure 5.A, left panel). These genes, along with others that exhibit similar functions from the Fig. 5A left panel, are represented in the bar graph to exhibit fold change indicating the differences in brain transcriptome results as a response to NMES in WT and Kl^HET^ (Figure 5.B) [30,33,34]. Conversely, downregulated genes such as *Copb2* indicate relation to vesicle formation in myoblasts as a result of treatment [33] (Figure 5.A, left panel). The volcano plot for Kl^HET^ NMES as compared to Kl^HET^ control indicates upregulation in 155 genes, including *Erg* which has been previously linked to angiogenesis upregulation, inhibition of vascular inflammation reduced response to immune stimuli, and downregulation in 200 genes including *Cd74* which promotes immunosuppression [36–39] (Figure 5.A, right panel). Using a dot plot to represent DAVID GO Terms from DEGs in Fig. 5A, we discovered little to no overlap between biological processes associated with the DEG in both genotypes as a result of NMES (Figure 5.C). NMES treatment in the WT group indicate upregulated GO Terms including blood vessel morphogenesis (*Edn1*), blood vessel diameter maintenance (*Avp*), and positive regulation of cytosolic calcium ion (*Oxt* and *Calcr*) while Kl^HET^ mice displayed establishment of epithelial cell polarity (*Cyth1*), apoptotic signaling by p53 mediator (*Wwox*), and mRNA processing (*Barx2*, *Lsm7*, and *Stk11*) (Figure 5.C, left panel). Downregulated GO Terms from WT mice include vesicle-mediated transport (*Copb2* and *Chmp1a*), DNA damage response (*Mgme1*), and ribosome biogenesis (*Ltv1*) while the Kl^HET^ comparison includes extracellular matrix organization (*Vwa1*), collagen fibril organization (*Col3a1*), and positive regulation of interleukin-6 product (*Cd74*), neurological pathway response to treatment may be genotype-specific (Figure 5.C, right panel).

**Figure 5.**
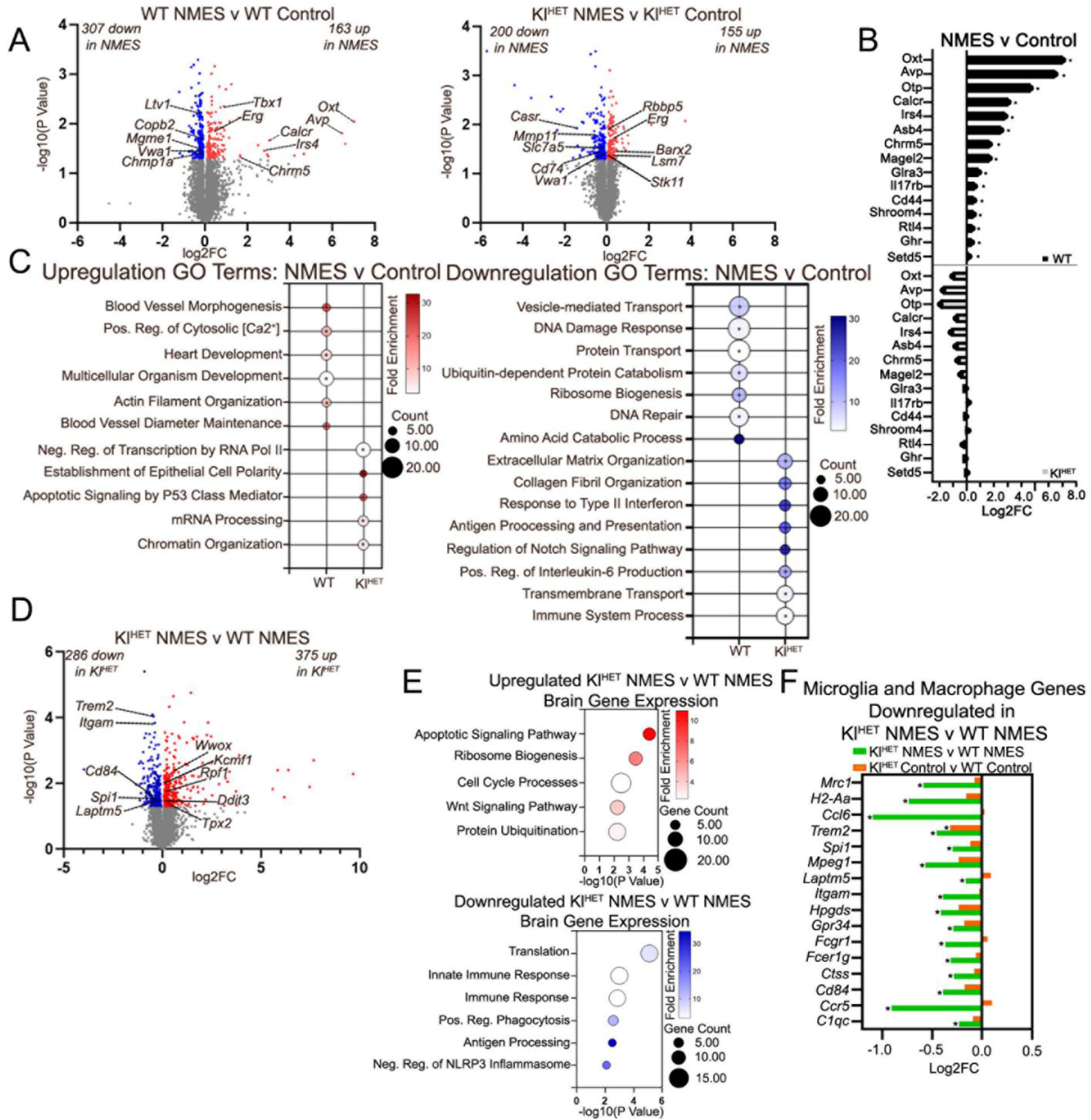
Effect of NMES on brain transcriptome indicates little overlap in WT compared to Kl^HET^. Gene expression profiling was performed by RNA-seq on dissected brain from WT and Kl^HET^ mice that underwent NMES treatment and edgeR was utilized to identify differentially expressed genes between treatment groups. **(A)** Volcano plots represent differentially expressed genes (DEGs) between NMES and control in WT mice (left panel) and in Kl^HET^ (right panel) at *p*<0.05. Red denotes significantly upregulated genes, blue for significantly downregulated, and grey denotes non-significance. **(B)** Bar graph represents selected DEGs from WT-NMES vs WT-control in both genotypes. **(C)** The bubble plot denotes top GO terms with the highest fold enrichment in each genotype from both upregulated (left) and downregulated (right) DEGs as generated by DAVID. **(D)** Volcano plots represent differentially expressed genes (DEGs) between Kl^HET^ NMES and WT NMES at *p*<0.05. Bubble plots depicting Gene Ontology (GO) terms for biological functions associated with DEGs shown on panel D was performed using DAVID. Upregulated terms are shown on the upper panel and downregulated terms on the lower panel. Stacked bar graph represents a comparison of fold change of selected microglia and macrophage DEGs downregulated in Kl^HET^ NMES v WT NMES. (N: WT 6/group; Kl^HET^ control=11; Kl^HET^ NMES=12). \**p*<0.05

To evaluate the interaction between genotype and exercise condition in the brain, we evaluated DEGs from Kl^HET^ NMES versus WT NMES with all genes shown on the volcano plot in Fig. 5D. The 375 genes upregulated in Kl^HET^ NMES versus WT NMES were represented by GO terms apoptotic signaling pathway (*Wwox*), ribosome biogenesis (*Rpf1*), cell cycle processes (*Cdc73* and *Ddit3*), Wnt signaling pathway (*Wwox*, *Cdc73*, and *Ddit3*), and protein ubiquitination (*Kcmf1*) (Fig 5.E, upper panel). Downregulated GO terms include translation (*Rpl3*), innate immune response (*Trem2* and *Cd84*), immune response (*Ccr5*), positive regulation of phagocytosis (*Fcrg1*, *Itgam*, and *Fcerg1*), and negative regulation of NLRP3 inflammosomes (*Trem2*). Notably, downregulated GO terms are predominantly related to immune function, found in macrophages and microglia. Therefore, evaluation of genes microglia- and macrophage-specific were represented by fold change comparing the treatment status of Kl^HET^ and WT (Figure 5.F). The only gene significant in both the NMES and control comparisons is *Trem2*, the deficiency of which is highly correlated with Tau aggregation and hyperphosphorylation and AD status [40]. The immune suppression, while a typical result of KLOTHO deficiency, is more pronounced in the NMES condition comparison between genotypes than the control [41]. This is likely due to the fact that temporary immune suppressant is common following exercise [42]. The difference in fold change is an indicator that the localized muscular stimulation has distal effects on the brain which is modified by the KLOTHO deficiency. Interestingly, the only gene which was upregulated by NMES in both genotypes is Erg, a transcription factor that impacts endothelial junction stability and angiogenesis [43]. The only shared downregulated gene is *Vwa1* which is involved in extracellular matrix organization and the mutation of which results in delayed neuromuscular responses and impaired motor coordination [44,45].

## Discussion

Within the experiment, we show that DEGs within stimulated muscle from NMES are related to metabolic functions and cellular structural modifications. Genes related to insulin regulation and metabolic modification such as *Lep*, *Nnat*, *Gck*, *Acvr1c*, *Retn*, *Ghr*, *Adipoq*, and *Rbp4* were differentially expressed in muscle tissue at the direct area of stimulation for all NMES condition compared to control indicating a metabolic pathway response similarity between genotypes in local tissue. For example, several processes, including cellular metabolism, energy balance, and inflammation reduction with LEP resistance leading to weight gain, cardiac inflammation, and hypertension [46]. The upregulation of *Lep* within our data indicates greater LEP production in direct areas of stimulation, thus proposing a metabolically favorable local effect of NMES treatment which could play a role in weight maintenance for both WT and KLOTHO-deficient organisms (Figure 2.G, left panel). Evaluation of previous work indicates *LEP*’s predominant role in obesity-related disruption of synaptic plasticity due to its strong link to obesity and its strong relationship to insulin-related genes [47,48]. Therefore, ncRNA analysis from plasma samples that indicate ncRNA which may regulate the trafficking of genes related to *Lep* such as from muscle to other areas of the body provides support that localized stimulation may impact the release of EV contents to distal organs (Figure 4.D). Despite some shared lipid and carbohydrate metabolism DEGs, most exercise-resultant localized DEGs were genotype stratified yet supporting similarity of biological functions related to glucose metabolism and insulin response (Figure 2.F-G). Therefore, our data supports the conclusion that similar biological results are achieved via predominantly different DEG pathways in localized tissue which a likely function of KLOTHO deficiency.

While the brain data indicates a response to NMES which supports the hypothesis that localized NMES has distal effects, a difference in both DEG function and composition are present. Only WT brain data shows enrichment of several genes related to metabolic function, cognition, and blood flow including *Oxt*, *Avp*, *Otp*, *Calcr*, *Irs4*, *Asb4*, *Chrm5*, *Magel2*, *Glra3*, *Il17rb*, *Cd44*, *Shroom4*, *Rtl4*, *Ghr*, and *Setd5* [29,49,50].

Notably, results of enriched *OXT* and *AVP* induce both insulin and glucagon for increased metabolic functioning, so to see a modification in brain data from local stimulation indicates promising neurological metabolic benefits from the localized activity [51,52]. Importantly, lipid metabolic benefits noted by *CD44* in brain which plays a role in lipid metabolic processes and insulin resistance, commonly associated with obesity, diabetes, and inflammatory disease [34,53]. Given that *Cd44* is downregulated, the role of insulin resistance is decreased, and lipid metabolism is increased to indicate that the metabolic pathway effects of NMES are consistent in whole brain tissue and muscle tissue in WT conditions. Importantly, none of these genes are noted as DEGs for Kl^HET^ NMES animals versus the controls which see very few genes related to glucose metabolic processes including *Pomc*, *Stk11*, *Gk*, and *Spop*. While previous work evaluates the role NMES has on muscle mass, heart rate, bone density and blood glucose, our recent findings provide specific genetic support to indicate how this treatment has widespread metabolic impact as exhibited by mRNA and ncRNA between KLOTHO-deficient and WT animals which is novel [54,55].

Notably, the microglia and macrophage gene data indicate significant differences between KLOTHO-deficient and WT treatment conditions related to immune function which is strongly tied to neurological disease (Figure 5.F) [56]. The significant downregulation of these immune-related genes in Kl^HET^ NMES versus WT NMES but not Kl^HET^ control versus WT control is consistent with previous exercise studies that note short-term exercise-induced immune stress as a means of strengthening the immune system in the long term [42]. Given sample collection less than 24 hours following the final round of treatment, the difference between Kl^HET^ and WT animals is likely indicative of the interplay between KLOTHO deficiency and localized exercise treatment on the brain. This result provides support for the argument that NMES differentially metabolically impacts KLOTHO-deficient models in the brain.

GO Term analysis of both muscle and brain mRNA indicated genetic modifications for processes involving structural elements like muscle sarcomere organization and blood vessel organization that indicates metabolic benefits to NMES treatment. Within the muscle data, there is an upregulation of sarcomere organization for all NMES mice which indicates muscular growth as a result of the exercise-induced contractions (Figure 2.C) [57]. Specifically, upregulated genes *MYBPH, MYH6*, and *MYOM2* which regulate myosin binding indicate that sarcomere re-organization from NMES increases myosin binding to allow muscle contractions to grow muscle strength and size [58]. Increased muscular strength related to fast-twitch muscles in the EDL have reported by previous NMES trials, so our results supporting a genetic modification mechanism for the structural change to achieve increased metabolic action [59]. Additionally, downregulation in both WT and Kl^HET^ conditions of angiogenesis which is associated with modifications in capillary regression and metabolic diseases like Type 2 Diabetes supports this finding [60,61]. Notably, comparing GO terms from each genotype as stratified by treatment condition, similar muscular structural benefits were present by genotype. Both treatment and control conditions for Kl^HET^ saw upregulation in broad structural modification like cell adhesion, axon guidance, cell migration, and extracellular matrix organization the NMES treatment specifically resulted in collagen fibril organization, regulation of cell shape, and actin cytoskeleton organization (Figure 3.F). These results indicate that benefits from NMES treatment likely modify muscle structural components differently given KLOTHO deficiency which is a novel finding. However, comprehensive histological analysis of muscle tissue was not performed within the scope of this experiment.

Genes related to brain vasculature differences emerged between genotypes given treatment comparison. Increased brain vasculature, especially in areas for memory consolidation as in hippocampal vascularization, was shown to improve cognitive performance [62]. Therefore, the speculative upregulation of blood vessel morphogenesis and blood vessel diameter maintenance from transcriptomic results in NMES treatment conditions as compared to controls signifies a potential blood flow benefit to WT conditions as a result of localized exercise (Figure 5.D). However, Kl^HET^ shows GO term downregulation of collagen fibril organization, and considering its regulatory role in maintaining vascular stiffness and strength, this transcriptomic response could be an indication of vasculature modification via NMES exercise (Figure 5.D) [63]. Therefore, NMES likely modifies brain vasculature via differing modalities in WT and Kl^HET^ animals. The structural differences could be explained by the previously established role of KLOTHO deficiency in cranial vascular calcification, necessitating different modalities for blood flow [64,65]. Similarity of certain functions with a difference in gene expression may be partially explained by the shared upregulated gene, (*Erg)*. Commonly known as a master transcription factor, *ERG* promotes transcription factor cascades regulating genes associated with blood flow and BBB structural maintenance via endothelial homeostasis [66].

To assess the contents of the transport mechanism between muscle and brain, we analyzed ncRNA signatures from plasma EVs. We chose to focus on miRNA data from this set due to little significance in the other types in response to NMES. However, analysis of circRNA data is included in supplemental results due to its high quantity of EV contents demonstrated in Fig 5A while simultaneously being related to unspecific processes i.e., phosphorylation, protein phosphorylation, and various protein metabolism and transports (Figure S.2D). Notably, miRNA significant in all NMES versus all control comparisons show similar directionality in both WT treatment comparisons and Kl^HET^ treatment comparisons (Figure 5.B). The biological responses such as response to oxygen levels and long-term synaptic potentiation are shared by treatment between genotypes (Figure 4.F). The only shared miRNA upregulated in each genotype group was let-7d-5p which belongs to the let-7 family of miRNAs which broadly regulate development [67]. More recent literature indicates that let-7d-5p presence regulates insulin signaling targets, correlated to increased obesity, and induces microglial release of inflammatory cytokines [68,69]. In previous human acute muscular exercise trials, the presence of let-7d-5p is upregulated and correlated with inflammatory responses [45]. Functional overlap in EV parallels that in muscle, providing support for the argument that localized muscular stimulation creates rejuvenation in the plasma. However, distal impacts of NMES across the BBB reveal KLOTHO-mediated functional differences. It is important to note that due to their size, EVs are difficult to isolate and once isolated, EVs vary in contents [70]. While we provide evidence that EVs were successfully isolated from plasma (Figure S.1), we did not perform genetic protein tagging for specific EV contents in vivo [71]. Tracking EVs from their release point in the muscle directly into the plasma and directly up to the choroid plexus without mediation from other organ systems would provide definitive proof for our proposed mechanism. However, these techniques are beyond the scope of our study.

Overall, the results of our study indicate that transcriptomic effects of localized muscular stimulation in brain are mediated by KLOTHO deficiency. While sharing similar biological functions in both muscle mRNA and plasma-derived EV ncRNA, the specific genes and ncRNAs responsible for the same biological processes varied by KLOTHO deficiency status. Brain mRNA data showed both differing biological function and gene expression indicating diverging biological processes as a consequence of communication with the stimulated area. Therefore, the cognitive transcriptomic impacts of localized stimulation diverged as a result of KLOTHO deficiency.

## Supporting information

Supplemental Images

---

**Supplemental Figure S1.**
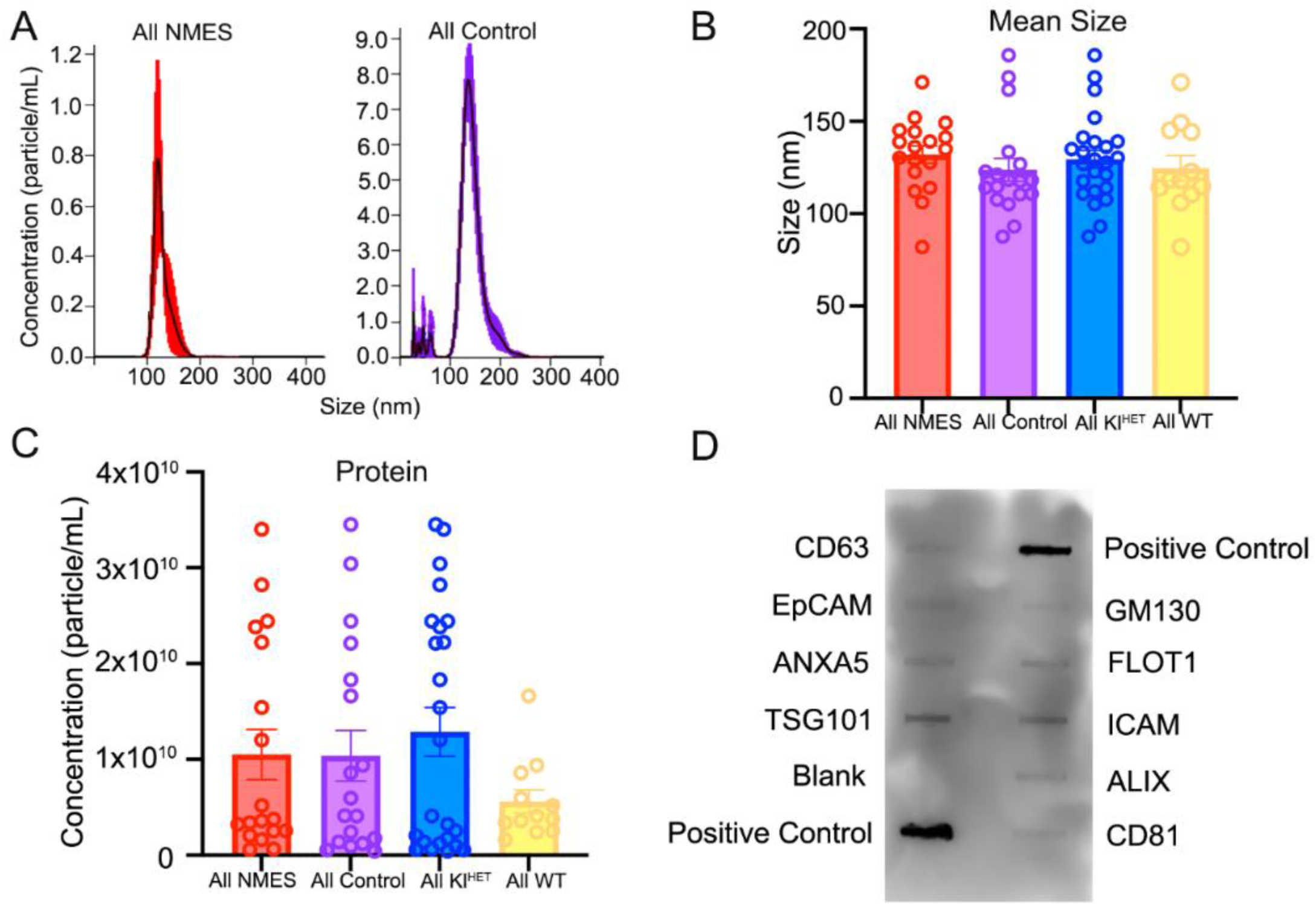
Plasma EV characterization indicates successful EV isolation and no significant size and concentration difference between genotypes or exercise conditions. Representative histogram **(A)** depicting particle size distribution determined by NanoSight-based nanoparticle tracking analysis of all NMES (A, left) and all control (A, right) plasma EVs. Bar plots summarizing mean size **(B)** or concentration **(C)** of isolated EVs. **(D)** Array confirming protein presence. (N: WT 6/group; Kl^HET^ control=11; Kl^HET^ NMES=12)

**Supplemental Figure S2.**
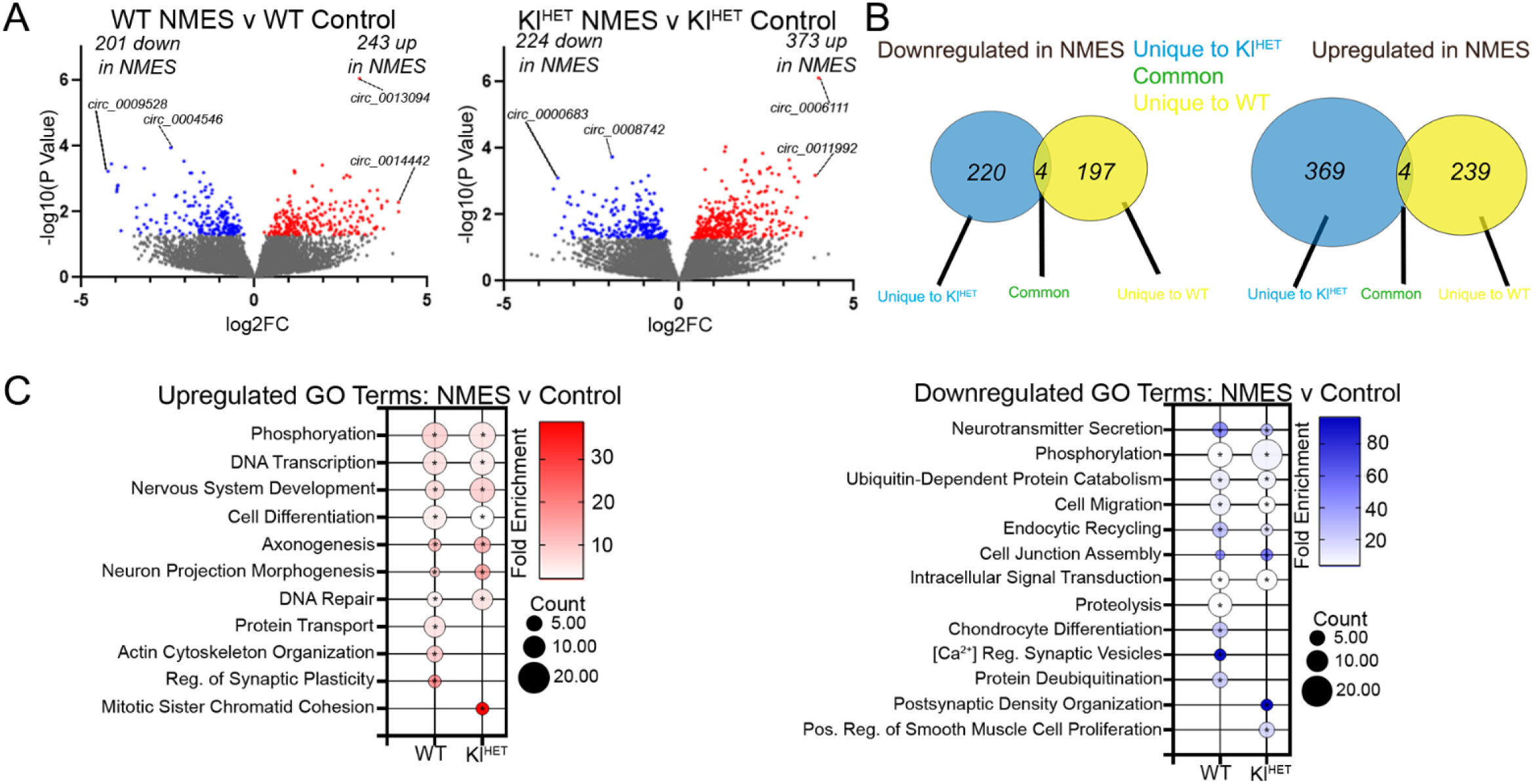
circRNA signatures plasma EVs associated with NMES in WT and Kl^HET^ mice. RNA was isolated from plasma EVs followed by small noncoding RNA sequencing to determine changes in EV cargos associated with experimental groups. **(A)** Volcano plots of differentially enriched circRNAs from NMES versus control comparisons in WT (left panel) and Kl^HET^ (right panel). **(B)** Venn diagram represents overlap in data from panel A (left and right) of circRNAs upregulated and downregulated in NMES v control. **(C)** Bubble plots depicting gene ontology (GO) terms for biological functions from data in panel A (left and right) using DAVID. Upregulated terms are shown on the left panel and downregulated terms are shown on the right panel. (N: WT 6/group; Kl^HET^ control=11; Kl^HET^ NMES=12). \**p*<0.05

## Institutional Review Board Statement

The animal study protocol was approved by the Institutional Review Board of University of Pittsburgh (PHS Assurance Number: D16-00118) on 10/18/2019 (Protocol #: 19105944).

## Data Availability Statement

The data that support the findings of this study were deposited to NCBI Gene Expression Omnibus (GEO) database and will be publicly available after the manuscript is accepted.

## Author Contributions

Conceptualization, R.K. and I.L., Methodology, C.A.C. and A.E.M. N.F.F; Investigation A.E.M.; Formal analysis C.A.C., A.E.M, R.K. Visualization C.A.C., A.E.M, Writing-original draft C.A.C., A.E.M. and R.K.; Writing-review & editing R.K., N.F.F., I.L..; Supervision R.K.; Project administration R.K.; Resources R.K. and I.L.; Funding acquisition R.K. and I.L. All authors have read and agreed to the published version of the manuscript.

## Materials and Methods

### Mouse Model

Animal studies were approved by the University of Pittsburgh Institutional Animal Care and Use Committee governed by regulatory standards from the Guide for the Care and Use of Laboratory Animals and the United States Department of Health and Human Services. Wild-type (WT) and KLOTHO HET (Kl^HET^) mice were internally bred for this experiment and randomly assigned to either the control group or NMES treatment group. The Kl^HET^ mouse strain used for this research project, B6;129S5-*Kl^tm1Lex^*/Mmucd, RRID:MMRRC_011732-UCD, was obtained from the Mutant Mouse Resource and Research Center (MMRRC) at University of California at Davis, an NIH-funded strain repository, and was donated to the MMRRC by Lexicon Genetics Incorporated. Mice were housed in a 75°C, 12:12 h light-dark cycled room with unrestricted access to standard food and water.

### NMES

Electrical stimulation was performed with a Neuromuscular Stimulator (Empi 300 PV, Boston, USA) at 9 mA under 2.5% isoflurane anesthesia. Mice were shaved on both right and left hind legs along the sciatic nerve and exposed skin was coated in O.C.T. compound to ensure proper probe placement and to prevent dermal burns. The double-pronged pen attachment was placed at piriformis, causing leg extension. NMES was delivered in sets of 10 consecutive stimulations per side, oscillating between each leg after each set, for a total of 2 sets on each leg. NMES mice received treatment every other day for a total of 5 treatment sessions. Perfusions and sample collection occurred 24 hours after the final treatment session. Control mice received 15 minutes of 2.5% isoflurane anesthesia exposure each day the NMES treatment group received stimulation.

### Sample Collection and Storage

Avertin-anesthetized mice were perfused with chilled-PBS after blood was collected intracardially with an EDTA-filled needle. Blood was centrifuged at 13,000 g for 5 minutes for separation of plasma and downstream processing. The grouping of tibialis anterior (TA) and extensor digitorum longus (EDL) muscles were collected from each leg, along with whole brain tissue. All samples were snap-frozen in dry ice before long-term storage in -80°C.

### Isolation of EVs from Plasma

Plasma samples were thawed at 4°C and 400 µL was processed with Norgen Biotek’s Plasma/Serum Exosome Purification kit (CAT# 57400), following manufacturer’s instructions. Aliquots of EV solution were kept for characterization and the remaining portion of solution was immediately used for RNA extraction.

### Extracellular Vesicle Characterization

All EV’s were characterized by Nanosight NTA software (v 3.4) to determine size and concentration. EV solution was diluted 1:500 and ran through tubing at 0.02 ml/min. EV protein markers were confirmed with Exo-Check Exosome Antibody Array (SBI EXORAY210B-8) following manufacturer instructions.

### RNA Extraction from EVs

Total RNA was isolated from the 190 µl elution of EV solution with Norgen Biotek’s Exosomal RNA Isolation Kit (CAT# 58000), following manufacturer’s instructions. Quality and concentration were monitored by the RNA Pico 600 Assay Kit for the Agilent 2100 Bioanalyzer System (5067-1513).

### Isolation of RNA from Brain

30 mg brain sections were subjected to RLT buffer from RNeasy Mini Kit (Qiagen 74106) with 10% beta-mercaptoethanol added and passed through a 25G needle. Homogenate was transferred to QIAshredder (Qiagen 79656) for further dissociation. The remaining manufacturers protocol was followed to complete the RNA isolation. Quality and concentration were analyzed via RNA Nano 600 kit for Agilent 2100 Bioanalyzer (5067-1511).

### Isolation of RNA from Muscle

50 mg muscle sections were powdered by the addition of N_2_ and manually dissociated via mortar and pestle. QIAzol (Qiagen 5,346,994) was added to the powdered muscle and passed through a 25G needle. Homogenate was added to QIAshredder and the RNeasy Mini Kit protocol was followed for the duration of RNA isolation. Quality and concentration were analyzed via RNA Nano 600 kit for Agilent 2100 Bioanalyzer (5067-1511).

### RNA Library Generation from EVs

Small non-coding libraries were produced by New England BioLabs’ NEBNext Multiplex Small RNA Library Prep Set (E7300S and E7580S). Library quality was assessed on samples by the High Sensitivity DNA Assay Kit for the Agilent 2100 Bioanalyzer System (5067-4626). Samples were sequenced by the UPMC Genome Center, Pittsburgh PA, on the NextSeq 2000 machine generating single-end reads. Data was received in .fastq.gz formatting.

### RNA Library from Brain and Muscle Tissue

Extracted RNA was sent to Novogene Co. located in Sacramento CA, for generation of total RNA libraries and sequencing on NovaSeq X Plus Series (PE150) machine. Data was received in .fq.gz formatting.

### RNA Alignment and Differential Enrichment

Brain and muscle mRNA data were aligned with Rsubread (v 2.16.1), annotated with the org.Mm.eg.db package (v 3.19.1) and analyzed with edgeR (v 4.0.16). Gene ontology (GO) terms were generated in DAVID with a p-value cut-off of <0.05 followed by a fold-change cut-off of >0.5 [72,73]. Noncoding RNA library data were aligned and quality checked by STAR (v 2.5.3a) and analyzed with COMPSRA (v 1.0.3) and DESeq2 (v 1.42.1). Gene ontology (GO) terms were generated in DAVID with a p-value cut-off of <0.05. Selected gene targets were analyzed using miRDB at target score cutoff>50. Representative gene visualization was created utilizing STRING interaction network database.

### Statistical Analysis

Sample sizes are indicated in the legends and correspond to biological replicates. Power analysis was performed to estimate the number of animals (two groups, t test, G*Power v3.1) with individual estimated experimental effect size, alpha =0.05, and 95% power. Figure legends include sample size (n) information for each treatment group. EV characterization was analyzed with a two-tailed unpaired t-test. Biological function terms were reduced by redundancy utilizing REVIGO, reduce+visualize Gene Ontology, (v 1.8.1). Unless otherwise stated, statistical analysis and representation was visualized in GraphPad Prism (v 10.0.3).

