## Supplemental Images for "Transcriptomic Response to Neuromuscular Electrical Stimulation in Muscle, Brain and Plasma EVs in WT and Klotho-deficient Mice"

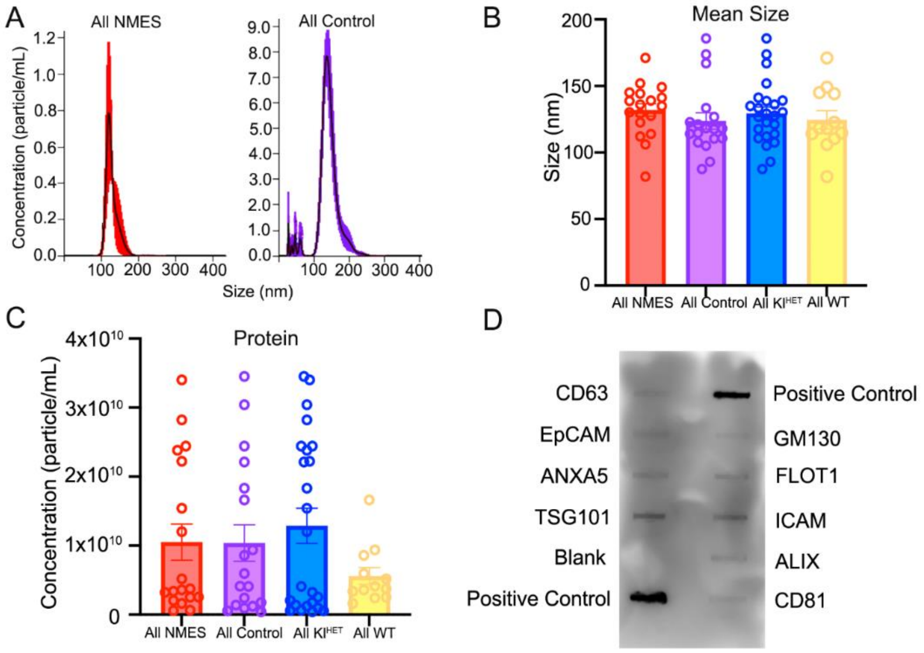


**Supplementary Figure S1: Plasma EV characterization indicates successful EV isolation and no significant size and concentration difference between genotypes or exercise conditions.** Representative histogram **(A)** depicting particle size distribution determined by NanoSight-based nanoparticle tracking analysis of all NMES (A, left) and all control (A, right) plasma EVs. Bar plots summarizing mean size **(B)** or concentration **(C)** of isolated EVs. **(D)** Western blot confirming protein presence. (N: WT 6/group; $KI^{HET}$ control=11; $KI^{HET}$ NMES=12)

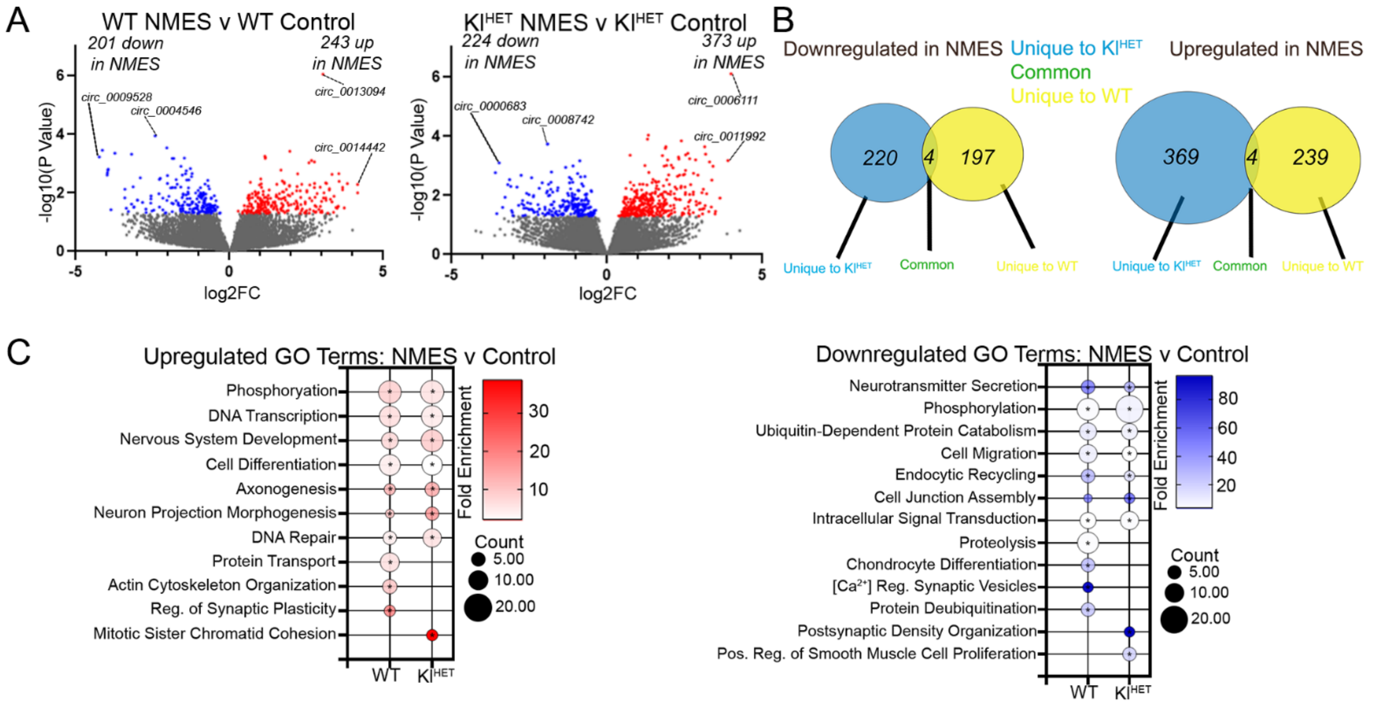


**Supplemental Figure S2. circRNA signatures plasma EVs associated with NMES in WT and KI$^{HET}$ mice**. RNA was isolated from plasma EVs followed by small noncoding RNA sequencing to determine changes in EV cargos associated with experimental groups. **(A)** Volcano plots of differentially enriched circRNAs from NMES versus control comparisons in WT (left panel) and KI$^{HET}$ (right panel). **(B)** Venn diagram represents overlap in data from panel A (left and right) of circRNAs upregulated and downregulated in NMES v control. **(C)** Bubble plots depicting gene ontology (GO) terms for biological functions from data in panel A (left and right) using DAVID. Upregulated terms are shown on the left panel and downregulated terms are shown on the right panel. (N: WT 6/group; KI$^{HET}$ control=11; KI$^{HET}$ NMES=12). *$p$<0.05
